# Histone methyltransferase MET-2 differentially regulates autosome and X-chromosome structure and transcription during meiosis in *C. elegans*

**DOI:** 10.64898/2026.07.26.740854

**Authors:** Carolyn Remsburg, Aimee Jaramillo-Lambert

## Abstract

During meiosis, accurate chromosome segregation requires significant condensation and compaction. These processes are mediated by condensins, cohesins, and histone tail modifications. We identified that MET-2, a histone methyltransferase that catalyzes the dimethylation of histone H3 lysine 9 (H3K9me2), differentially impacts chromosome size in the male vs. female *C. elegans* germline. In *met-2* null worms, autosomes during spermatogenesis are significantly larger than wild type, while chromosome size during oogenesis is unaffected. X-univalent size in males is also unaffected by loss of MET-2, indicating MET-2 differentially regulates autosomal and X-chromosome compaction in male spermatogenesis. Autosome size is not changed when males harbor a catalytically deficient MET-2 *(met-2*CD) or have mutations preventing germline histone H3K9 methylation (H3K9R). In addition, *met-2* males, in contrast to *met-2*CD or H3K9R males, have more active RNA pol II in later stages of meiosis. These data suggest MET-2 plays a noncatalytic role in mediating chromosome structure and transcription. In *met-2* male germ lines, genes on the X chromosome, which is typically enriched in H3K9me2, are significantly more likely to be upregulated than genes on autosomes, even though X-univalent size is unchanged. These results suggest that MET-2 plays a sex-specific role that is not limited to its enzymatic activity.

## Introduction

Chromosome compaction must be highly regulated during meiosis and mitosis, as improper compaction leads to chromosome segregation defects (Yu and Koshland 2003; Kagami and Yoshida 2016). Several protein complexes regulate chromosome compaction, such as cohesins and condensins (Csankovszki et al. 2009; Ravi et al. 2020; Kim and Yu 2020). In addition, histone post-translational modifications (PTMs) are known to regulate chromosome compaction (Rea et al. 2000). Dimethylation of lysine 9 of histone 3 (H3K9me2), is especially associated with chromatin compaction (Padeken et al. 2022; Schultz et al. 2002).

During meiosis, there are sex-specific differences in chromosome compaction, such as the replacement of histones with protamines and protamine-like proteins to hypercompact DNA in spermatogenesis (Braun 2001; Shakes et al. 2009). In *C. elegans*, X-chromosomes have differences in histone methylation in oogenesis vs spermatogenesis (Kelly et al. 2002). In addition, MET-2 (SETDB1 in mammals), the histone methyltransferase responsible for H3K9me2, differentially impacts oogenesis and spermatogenesis in the presence of unsynapsed chromosomes. While MET-2 deposits H3K9me2 marks on unsynapsed chromosomes in both spermatogenesis and oogenesis, it only mediates checkpoint inactivation during spermatogenesis (Bessler et al. 2010; Checchi and Engebrecht 2011). Additionally, chromosomes enriched by MET-2 mediated H3K9me2 appear to be more compact during spermatogenesis compared to oogenesis (Checchi and Engebrecht 2011). In mammals, SETDB1 also plays a sex-specific role in regulating chromosome pairing and synapsis during spermatogenesis but does not appear to impact chromosome structure during oogenesis (Eymery et al. 2016b; Cheng et al. 2021).

Histone methylation is also correlated with transcriptional inhibition, the regulation of which may be context-specific (Bannister et al. 2001; Lachner et al. 2001). For example, in *C. elegans* embryos loss of MET-2 results in an overall derepression of transcription (Delaney et al. 2022) and in spermatogenesis, the loss of all H3K9 methylation causes global derepression of transcription (Chien and Michael 2023). However, in the hermaphrodite germline, derepression is not observed (Guo et al. 2015). In mammals, specific families of endogenous retroviruses are transcriptionally downregulated by SETDB1 in adult lineage-restricted B-cells, but not in embryonic stem cells, suggesting this context-dependence is evolutionarily conserved (Collins et al. 2015).

While the canonical role of MET-2 is to catalyze histone methylation, there is increasing evidence that MET-2 may have additional capabilities. Catalytically-deficient MET-2 still appears to repress transcription of many genes (Delaney et al. 2022). Non-catalytic roles for histone methyltransferases in regulating transcription have also been proposed in other species (e.g., mammalian G9a and GLP) (Tachibana et al. 2008). While a connection has been posited between transcription and chromatin compaction, the non-catalytic role of histone methyltransferases in regulating chromatin compaction during meiosis has not been explored.

Here, we probed the role of MET-2 and H3K9 dimethylation in regulating chromatin compaction and transcriptional repression during spermatogenesis. We found that MET-2 has a sex-specific role in regulating chromosome compaction. MET-2 also plays a non-catalytic role in regulating both chromosome compaction and global transcription. In addition, genes on the X-chromosome are more likely to be repressed by MET-2 than genes on autosomes. Overall, these results uncovered a noncatalytic role of MET-2 during spermatogenesis.

## Results

### MET-2 differentially regulates chromosome size in males versus hermaphrodites

To understand if histone methylation has a sexually dimorphic role in regulating meiotic chromosome size, we examined worms harboring a *met-2* deletion allele, *met-2(n4256)*, [referred to as *met-2(Δ)* throughout]. *met-2(Δ)* was previously shown to regulate germline H3K9me2 (Bessler et al. 2010; Delaney et al. 2022). We measured bivalent volume and length in *him-8* (control) and *met-2(Δ); him-8* adult hermaphrodites and males during diakinesis of meiotic prophase, when homologous chromosomes are at their most compact state. During oogenesis in hermaphrodites, we observed no significant difference in diakinesis bivalent volume or length between *him-8* and *met-2(Δ); him-8* worms (**Fig. 1a** and **Fig. S1a**). However, during spermatogenesis in males, we observed that bivalents are significantly larger and longer in *met-2(Δ); him-8* compared to *him-8* (**Fig. 1b** and **Fig. S1b**). This suggests that MET-2 plays a sexually dimorphic role in regulating chromosome compaction during diakinesis.

**Figure 1:**
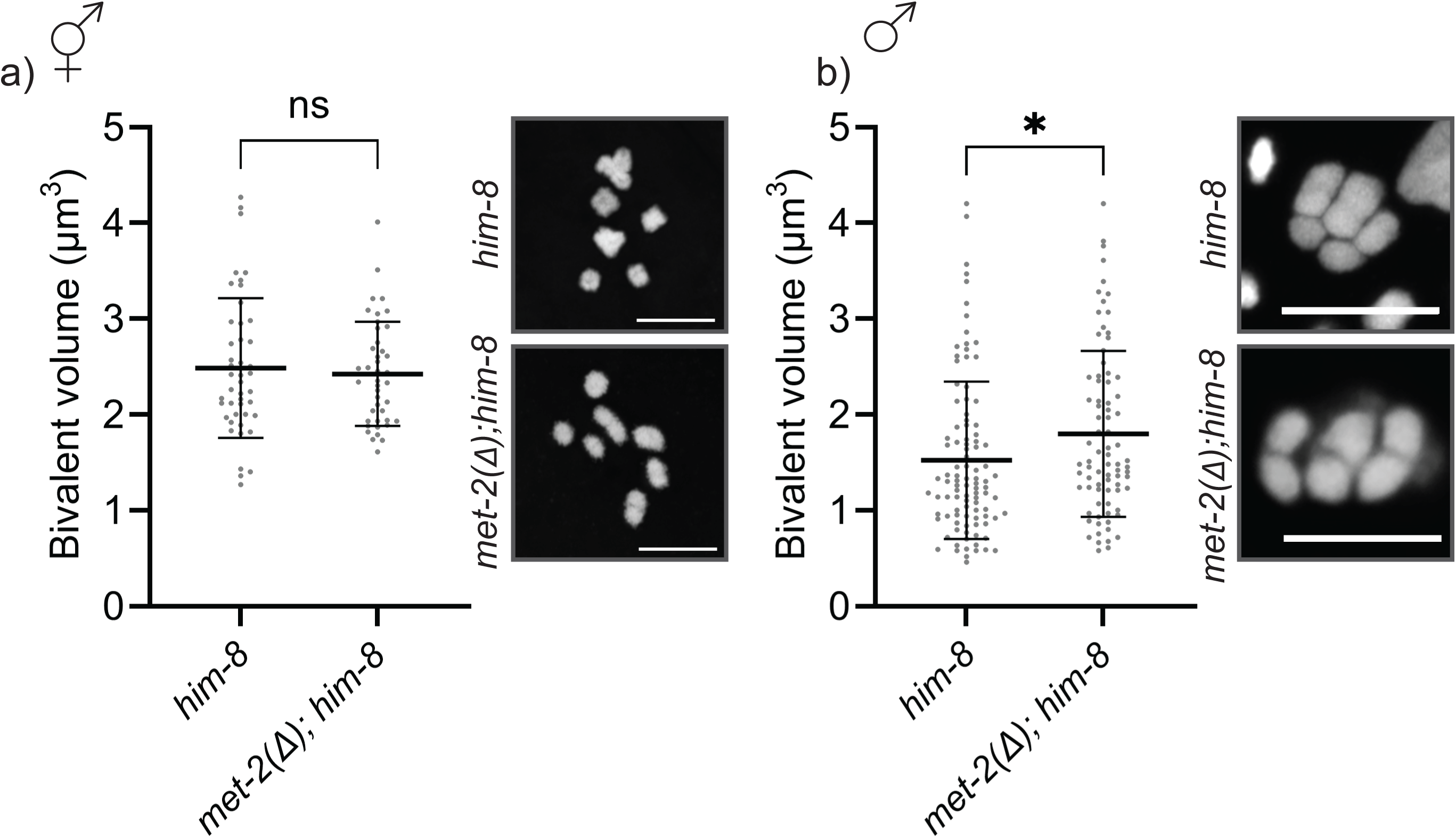
MET-2 differentially regulates bivalent volume in hermaphrodite oogenesis vs male spermatogenesis. Volume of individual a) hermaphrodite (N=45 *him-8*, 42 *met-2(Δ); him-8*) and b) male (N=100 *him-8*, 80 *met-2(Δ); him-8*) diakinesis bivalents in adult worms of the indicated genotypes. Bars represent mean bivalent volume, and error bars represent SD. ** p<0.01, ns = no significance using a t-test. DAPI stained images of a) hermaphrodite and b) male diakinesis nuclei of indicated genotypes. Scale bar = 5 µm.

### Diakinesis bivalent size is not affected in the absence of H3K9 dimethylation

Previous work suggested that MET-2 has additional functions beyond its role as a histone methyltransferase (Delaney et al. 2022). To determine if the increased diakinesis bivalent size observed in *met-2(Δ)* males is due to the catalytic activity of MET-2, we examined worms harboring a point mutation in the SET domain, which is the site of methyltransferase activity (Delaney et al. 2022). These *met-2(gw1660)* worms (referred to as *met-2*CD) were shown to have a similar reduction in H3K9me2 to *met-2(Δ)* worms, but have similar protein stability to wild-type (WT) MET-2 (Delaney et al. 2022). We measured diakinesis bivalents in males and found that there is no difference in volume or length in *met-2CD; him-8* males compared to *him-8* males (**Fig. 2a** and **Fig. S2a**).

**Figure 2:**
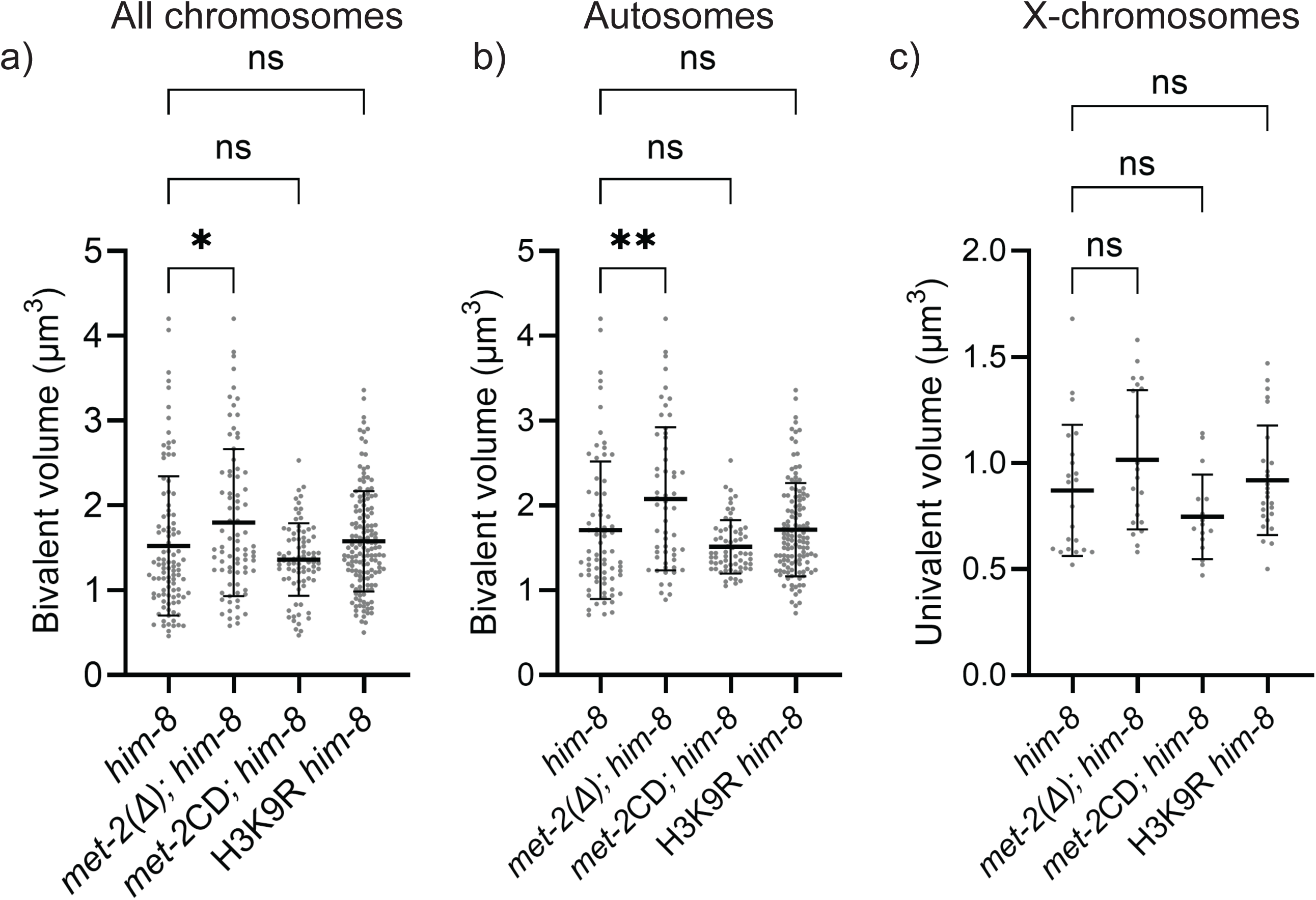
MET-2 regulates volume of the autosome independently of its catalytic activity. a) Volume of male diakinesis chromosomes of indicated genotypes.(N=100 *him-8*, 80 *met-2(Δ); him-8*, 81 *met-2*CD*; him-8*, 150 H3K9R *him-8*). b) Volume of male diakinesis autosomes of indicated genotypes. (N=73 *him-8*, 57 *met-2(Δ); him-8*, 65 *met-2*CD*; him-8*, 124 H3K9R *him-8*). c) Volume of male diakinesis X-chromosomes of indicated genotypes (N=22 *him-8*, 21 *met-2(Δ); him-8*, 16 *met-2*CD*; him-8*, 26 H3K9R *him-8*). Bars represent mean volume, and error bars represent SD. * p<0.05, ** p<0.01, ns = no significance using ANOVA with post-hoc Tukey-Kramer.

To understand if post-translational modifications of H3K9 directly regulate chromosome compaction, we created a strain harboring a mutation of lysine 9 (K9) of all histone H3s expressed in the germ line (*his-45*, *his-55*, *his-59* and *his-63*) (Gleason et al. 2023). Using CRISPR/Cas9 genome editing, we generated a strain in which all four germ line expressed histone H3s contain a point mutation which converts K9 to an arginine (H3K9R). We measured bivalent size during diakinesis in *him-8* and H3K9R *him-8* males. Similar to the *met-2*CD males, there is no difference in bivalent size in H3K9R *him-8* males compared to *him-8* males (**Fig. 2a** and **Fig. S2a**). These data suggest that MET-2 is playing a non-catalytic role in regulating chromosome compaction in spermatogenesis.

### X-chromosome compaction is not regulated by H3K9me2

H3K9me2 is enriched on unsynapsed chromosomes during meiosis and is highly enriched on the single X-chromosome in WT males (Bessler et al. 2010). Based on this observation, we tested if X-chromosome compaction specifically was affected when H3K9me2 was disrupted. To measure the X-chromosome, we immunolabeled for H3K4me2, a post-translational modification that is enriched on the autosomes during diakinesis (Samson et al. 2014). Interestingly, in contrast to the autosomes (**Fig. 2b** and **Fig. S2b**) we found no significant difference in the size of the X-chromosome in *him-8*, *met-2(Δ); him-8*, *met-2*CD*; him-8* or H3K9R *him-8* males (**Fig. 2c** and **Fig. S2c**). This shows that X-chromosome compaction in males is not directly regulated by H3K9me2.

### Global transcription during spermatogenesis is mediated by MET-2, rather than H3K9me2

As chromosome compaction has been associated with transcriptional inhibition, we next examined global transcription during meiosis when H3K9me2 is perturbed. To assess global transcription, we immunolabeled with an antibody that recognizes a PTM of RNA polymerase II associated with transcription elongation (RNA pol II pSer2), which is abruptly down-regulated at the karyosome stage in spermatogenesis (Shakes et al. 2009). We found that *met-2(Δ); him-8* males have a delayed inhibition of transcription compared to *him-8* males (**Fig. 3a, b**). This is consistent with previous literature (Chien and Michael 2023) and with the observation that these males have larger chromosomes (**Fig. 1b**). We also observed that *met-2CD; him-8* males have no change in transcriptional inhibition compared to *him-8* males (**Fig. 3a, b**), consistent with the lack of difference in bivalent size (**Fig. 2a**). Finally, we observed that H3K9R *him-8* males exhibit precocious inhibition of transcription compared with *him-8* males (**Fig. 3a, b**). As the mutation in H3K9R males should prevent all H3K9 PTMs, not just dimethylation, this suggests that H3K9 PTMs may contribute to both activation and inhibition of transcription.

**Figure 3:**
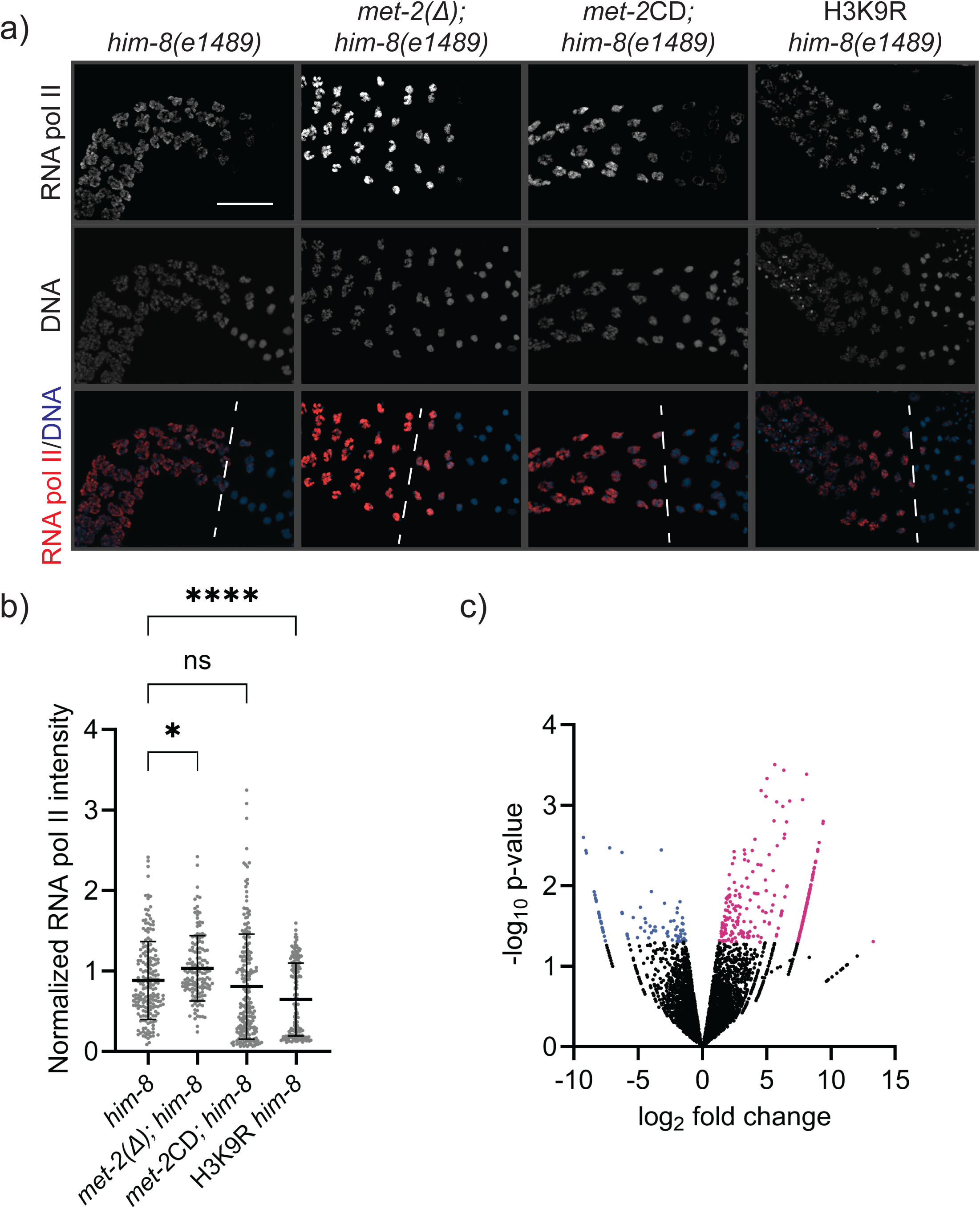
MET-2, but not H3K9me2 regulates global transcription in males. a) Male gonads of the indicated genotype were dissected and immunolabeled for active RNA polymerase II (RNA pol II Pser2). The white dotted line indicates the diplotene-karyosome transition. Scale bar = 20µm. b) RNA pol II intensity of late prophase I nuclei was normalized to pachytene nuclei and plotted. Bars represent mean ratio and error bars represent SD. * p<0.05, **** p<0.001, ns = no significance using ANOVA. (N=15 gonads for each genotype, 190 *him-8* nuclei, 143 *met-2(Δ); him-8* nuclei, 208 *met-2*CD*; him-8* nuclei, 180 H3K9R *him-8* nuclei). c) Volcano plot of genes differentially expressed in *met-2(Δ); him-8* vs *him-8* male gonads identified by RNASeq. The x-axis shows the log_2_ fold change and the y-axis displays the -log_10_ adjusted p-value. The threshold for significance is set at an adjusted p-value of <0.05. Significantly up-regulated genes are shown in magenta and significantly down-regulated genes are shown in blue. Unchanged genes are shown in black.

### X-chromosome genes are more likely to be transcriptionally regulated by MET-2

As we observed a global increase in transcription in *met-2(Δ); him-8* males, we were interested in understanding the specific transcripts exhibiting increased expression. We performed RNA-seq on gonads dissected from *him-8* and *met-2(Δ); him-8* males. We found 261 genes that were significantly upregulated in *met-2(Δ); him-8* males and 84 genes that were significantly downregulated (**Fig. 3c**), consistent with the idea that MET-2 downregulates transcription. No GO term enrichment was found among the differentially-expressed genes. In males, the X-chromosome is highly enriched in H3K9me2 and transcriptionally down-regulated (Kelly et al. 2002). We found that genes on the X-chromosome were more likely to be upregulated in *met-2(Δ); him-8* males compared to genes on the autosomes (**Fig. 4a, b**). In addition, genes on the X-chromosome were 3.0 times as likely as genes on the autosomes to be found among the 20 most highly upregulated genes (**Fig. 4c**). Using qPCR, we were able to validate the upregulation of a selection of genes on the X-chromosome (**Fig. S3**).

**Figure 4:**
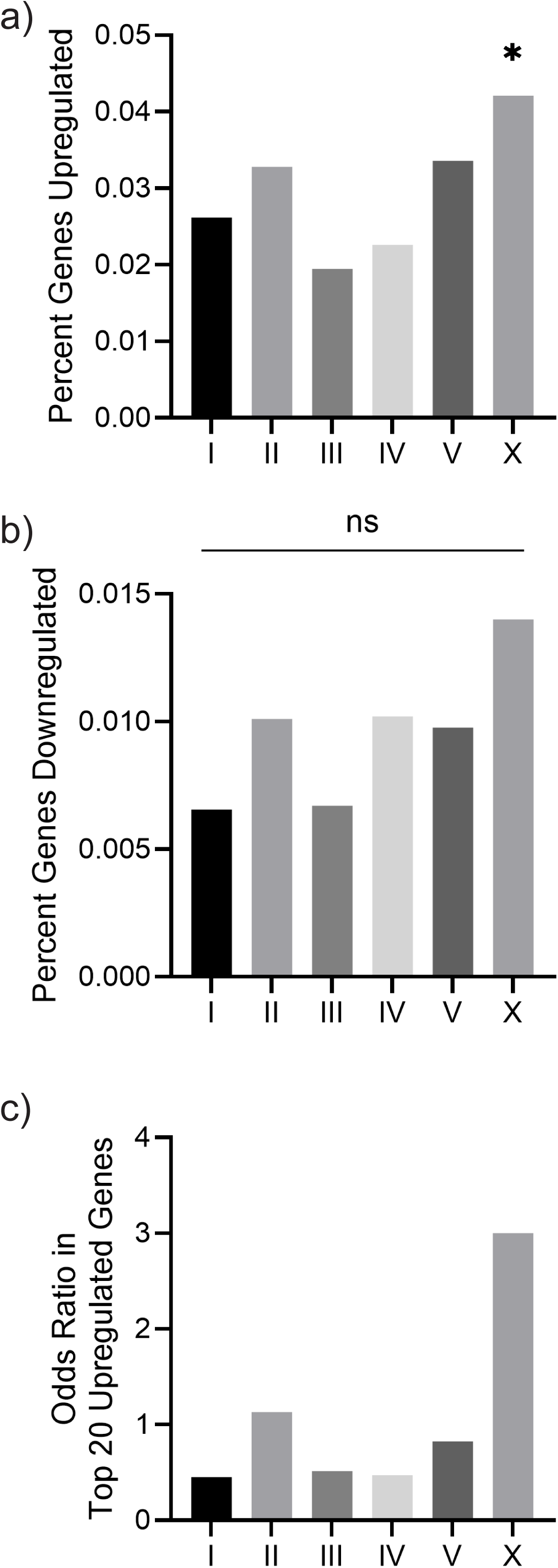
Genes on the X-chromosome are more likely to be upregulated in *met-2(Δ)* male germlines compared to autosomal genes. a) Genes on the X-chromosome were more likely to be upregulated compared to genes on the autosomes. b) No chromosome was overrepresented in the downregulated genes. c) Genes on the X-chromosome were 3.0 times as likely to be found in the twenty most highly upregulated genes. To test if more genes were upregulated on the X-chromosome compared to genes on the autosomes, the number of upregulated genes on each chromosome was compared individually to all the other chromosomes combined. Only the X-chromosome showed statistical significance.* p<0.05, ns=not significant using chi-square test.

## Discussion

Histone methylation, particularly at H3K9, is an important regulator of genome architecture and gene expression. In this work, we found that MET-2, a histone methyltransferase, plays a sex-specific, non-catalytic role in regulating both chromatin compaction and transcription during meiosis. Additionally, we found that MET-2 preferentially inhibits transcription of genes on the X-chromosome compared to genes on the autosomes.

A sex-specific role for MET-2 in regulating chromosome compaction during meiosis (**Fig. 1**) is consistent with previous studies examining histone methylation and chromosome structure during meiosis in other organisms. During spermatogenesis in mammals, flies and worms, the sex chromosomes are highly enriched in histone methylation catalyzed by SETDB1 (Alavattam et al. 2022; Hennig and Weyrich 2013; Bessler et al. 2010). In mice, SETDB1 is required for chromosome pairing and synapsis and loss of SETDB1 results in arrest at the zygotene stage (Cheng et al. 2021). However, during oogenesis, loss of SETDB1 does not affect folliculogenesis or chromatin structure (Eymery et al. 2016b; Kim et al. 2016). This differential effect of SETDB1 loss, along with our data supports the idea that SETDB1 has sex-specific roles during meiosis.

We found that MET-2 plays a non-catalytic role in regulating both chromosome compaction and transcription (**Fig. 2a, b and 3a, b**). This is consistent with previous work in embryogenesis where *met-2(Δ)* embryos have significantly more transcriptionally up-regulated genes than *met-2*CD embryos (Delaney et al. 2022). Together these data suggest that MET-2 plays additional roles, outside its methyltransferase activity. Perhaps this noncatalytic activity manifests through interactions with additional chromatin modifiers. One likely candidate is a histone deacetylase (HDAC), as embryos lacking MET-2 have increased H3K4 and H3K27 acetylation (Delaney et al. 2022). Additionally, histone methyltransferases in mammals and *Drosophila* are known to directly interact with HDACs (Yang et al. 2003; Czermin et al. 2001; Vaute et al. 2002).

During male spermatogenesis, the single X-chromosome is highly compacted and enriched in H3K9me2 catalyzed by MET-2 (Bessler et al. 2010). Interestingly, our results suggest that MET-2 does not regulate X-chromosome compaction but does regulate transcription of genes on the X-chromosome (**Fig. 2c and Fig. 4**). Previous work in *him-8(e1489)* hermaphrodite germ lines, which results in unsynapsed X-chromosomes enriched in H3K9me2, found that deleting MET-2 does not result in increased transcription of the X-chromosome (Guo et al. 2015). This contrasts with our results in males, where we found that in *met-2(Δ)* males, genes on the X-chromosome are more likely to be upregulated compared to genes on the autosomes (**Fig. 5**). This may indicate that sex-specific differences play a role in regulating X-chromosome transcription, or it may be due to a difference in X-chromosome number, as previous work showed MET-2 mediated H3K9 methylation inhibits transcriptional activation of the X-chromosome only when it is unpaired (Checchi and Engebrecht 2011).

To differentiate between the role of MET-2 and H3K9me2 in meiosis, we generated a strain in which the germline H3s all harbor a mutation at K9. We found that the H3K9R males phenocopied the *met-2*CD males in terms of bivalent size (**Fig. 2a, b**), supporting the hypothesis that MET-2 plays a non-catalytic role in regulating chromosome compaction. However, the H3K9R male germ lines exhibited a precocious inhibition of transcription (**Fig 3a, b**). As the H3K9R mutation prevents all PTMs of H3K9, including tri-methylation and acetylation, this is consistent with H3K9 acetylation promoting transcription (Gates et al. 2017).

Overall, this work indicates that MET-2 plays a sex-specific, non-catalytic role in regulating both chromosome compaction and transcription. Further experiments will help identify not only if MET-2 is acting as a scaffold for additional chromatin modifiers, but also how MET-2 regulates chromatin in a sex-specific manner.

## Materials and methods

### C. elegans Strains

All *C. elegans* strains used in this study were maintained on Modified Youngren’s, Only Bacto-peptone (MYOB) plates seeded with *E. coli* OP50 using standard culturing conditions (Brenner 1974). The following strains were utilized: N2, CB1489 *him-8(e1489)* IV, AJL139 *met-2(n4256)/qC1* III; *him-8(e1489)* IV, AJL132 *met-2(gw1660)* III; *him-8(e1489)* IV [*met-2*CD; *him-8*], AJL155 *his-45(ude57) his-55(ude58) his-59(ude59) his-63(ude60)* IV [H3K9R], AJL181 *his-45(ude57) his-55(ude58) his-59(ude59) him-8(ude67) his-63(ude60)* IV [H3K9R *him-8*]. Homozygous *met-2(n4256)*; *him-8(e1489)* [*met-2(Δ); him-8*] worms were maintained for less than 8 generations prior to being used for experiments.

### CRISPR/Cas-9 mediated genome editing

CRISPR/Cas-9-mediated genome edits were conducted using the clone-free homology directed repair method with *dpy-10* as a co-CRISPR marker (Paix et al. 2015; Arribere et al. 2014). Injections were conducted using an injection mix of 1.53 µM Cas9 protein (IDT, #1081058, Coralville, IA), 6.4 µM universal tracrRNA (IDT, #1072533, Coralville,IA), 1.25 µM *dpy-10* crRNA, 5 µM allele specific crRNA, 0.92 µM *dpy-10* repair oligonucleotide and 2.2 µM allele specific oligonucleotide. **Supplementary Table 1** lists crRNA and oligonucleotides. The edits were verified by Sanger sequencing and the presence of male progeny in the AJL181 strain. Primers are listed in **Supplementary Table S2**.

### Dissections and immunostaining

Dissections and immunostaining were performed as previously described (Rourke and Jaramillo-Lambert 2022). Briefly, hermaphrodites and males were synchronized by picking L4s and incubating at 20°C overnight. Gonads were dissected on a coverslip in 30 µL Egg Buffer (118 mM NaCl, 48 mM KCl, 2mM CaCl_2_, 2mM MgCl_2_, 25 mM HEPES, pH 7.4) with 0.1% Tween-20. Gonads were fixed by removing 15 µL liquid and replacing with 2% paraformaldehyde in Egg Buffer with 0.1% Tween-20. Another 15 µL of liquid was removed and the dissected gonads were covered with a Superfrost Plus slide. The gonads were fixed at room temperature for 5 min then submerged in liquid nitrogen. After a minimum of 5 min, the coverslip was rapidly removed, and the slides were immediately placed in -20°C methanol for 1 min. After washing 3 times for 5 min each in PBST (1XPBS, 0.1% Tween-20), the slides were blocked in 0.7% BSA in PBST for 1 h at room temperature. 50 µL of primary antibody diluted in 0.7% BSA in PBST was dispensed onto the gonads, and the slides were incubated at room temperature in a humid chamber overnight. Slides were washed 3 times for 5 min each in PBST. 50 µL DAPI (2 µg/mL) was dispensed on top of the gonads and the slides were incubated at room temperature for 5 min. The slides were washed 3 times for 5 min and gonads were mounted in VectaShield (Vector Laboratories, H-1000, Newark, CA). Antibodies and dilutions are listed in **Supplementary Table S3**.

### Imaging and post-image processing

Images were obtained with a Zeiss LSM880 Airy Scan confocal microscope (Carl Zeiss, Inc., Gottingen, Germany) using Zen software. Nuclei images were obtained using Z-steps of 0.1 µm and the full focal range was imaged. After AiryScan processing with Zen software, the images were deconvolved with Huygens Essential (Scientific Volume Imaging, Hilversum, The Netherlands). Imaris Image Analysis software (Oxford Instruments, Oxon, UK) was used to measure volume and length of individual bivalents. RNA pol II intensity was measured using ImageJ (Schneider et al. 2012). Individual nuclei were circumscribed using the freehand tool in the DNA track. The region of interest was opened in the RNA pol II track and the mean intensity was measured. For each gonad, five pachytene nuclei were averaged, then each individual nuclei in the condensation zone were normalized to that value. Three biological replicates were performed, consisting of five gonads per genotype per biological replicate.

### RNASeq

Gonad dissections for RNASeq was performed as previously described (Guo et al. 2015). *him-8* and *met-2; him-8* male L4s were staged overnight at 20°C. Adults dissected in “red PBS” (1X PBS, 10µM aurin tricarboxylic acid) and gonads pipetted into a microfuge tube containing 25 µL red PBS. Once 20-50 gonads were collected, 250 µL Nucleozol (Macherey-Nagel, #740404, Allentown, PA) was added to the tube, and stored at -80°C. After collection of 400 gonads of each genotype, samples were pooled and RNA was extracted using a Nucleospin kit for Nucleozol (Macherey-Nagel, #740406, Allentown, PA), purified using Nucleospin RNA Clean-up kit (Macherey-Nagel, #740948, Allentown, PA) and eluted in 30 µL RNase-free water. RNA was submitted for Ultra-Low Input RNA-Seq to Genewiz (Azenta Life Sciences, South Plainfield, NJ). Test statistics were done using a Wald test. Differential expression analysis data, including p-values, are listed in **Supplementary Table S4**. **Supplementary Table S5** contains raw counts. **Supplementary Table S6** lists normalized counts.

### qPCR

For each biological replicate of qPCR, 100 *him-8(e1489)* and *met-2(n4256); him-8(e1489)* males were picked into 25 µL “red PBS” and 250 µL Nucleozol was added. The samples were stored at -80°C until a total of 500 worms were obtained for each genotype. RNA was extracted as above and eluted in 60 µL RNase-free water. cDNA was synthesized using the iScript cDNA synthesis kit (BioRad, #1708891, Hercules, CA). qPCR was performed using PowerUp SYBR Green Master Mix (Thermo Fisher Scientific, A25742, Waltham, MA). Reactions were run on a QuantStudio 6 Real-Time PCR cycler system (Thermo Fisher Scientific, Waltham, MA), using manufacturer’s recommended settings. Threshold cycle (Ct) values were normalized first to *csq-1* and are depicted as the fold-increase of transcripts in *met-2(n4256); him-8(e1489)* compared to *him-8(e1489)* males. Primers were designed using Primer3 (Untergasser et al. 2012) (**Supplementary Table S2**).Three technical replicates per biological replicate were performed. Three biological replicates were performed.

### Statistical analysis

GraphPad Prism (Dotmatics, Boston, MA) was used to perform statistical analysis and graphing.

## Acknowledgements

Thank you to Susan Gasser and Jan Padeken for the MET-2CD strain. Thank you to Lauren Salvitti and Lacole Fung for the initial *met-2(Δ)* characterization and to members of the AJL lab for feedback on this project and manuscript.

## Data Availability

The authors state that all data necessary for confirming the conclusions are presented in the article, figures, and tables. *C. elegans* strains generated in this study are available upon request. Supplemental materials are available at *Genetics* online. RNASeq data are deposited at GEO (accession number pending).

